# *Drosophila* immunity: The *Drosocin* gene encodes two host defence peptides with pathogen-specific roles

**DOI:** 10.1101/2022.04.21.489012

**Authors:** M.A. Hanson, S. Kondo, B. Lemaitre

## Abstract

Antimicrobial peptides (AMPs) are key players in innate defence against infection in plants and animals. In *Drosophila*, many host defence peptides are produced downstream of the Toll and Imd NF-κB pathways. Use of single and compound AMP mutations in *Drosophila* has revealed that AMPs can additively or synergistically contribute to combat pathogens in vivo. However, these studies also revealed a high degree of specificity, wherein just one AMP can play a major role in combatting a specific pathogen. We recently uncovered a specific importance of the antibacterial peptide *Drosocin* for defence against *Enterobacter cloacae*. Here, we show that the *Drosocin* locus *(CG10816)* is more complex than previously described. In addition to its namesake peptide “Drosocin”, it encodes a second peptide generated from a precursor via furin cleavage. We name this peptide “Buletin”, and show that it corresponds to the uncharacterized “Immune-induced Molecule 7” previously identified by MALDI-TOF. The existence of a naturally occurring polymorphism (Thr52Ala) in the CG10816 precursor protein masked the identification of this peptide previously. Using mutations differently affecting the production of these two *CG10816* gene products, we show that Drosocin, but not Buletin, contributes to the *CG10816*-mediated defence against *E. cloacae*. Strikingly, we observed that Buletin, but not Drosocin, contributes to the *CG10816*-mediated defence against *Providencia burhodogranariea*. Moreover, the Thr52Ala polymorphism in Buletin affects survival to *P. burhodogranariea*, wherein the Alanine allele confers better defence than the Threonine allele. However, we found no activity of Buletin against either *P. burhodogranariea* or *E. coli* in vitro. Collectively, our study reveals that *CG10816* encodes not one but two prominent host defence peptides with different specificity against different pathogens. This finding emphasizes the complexity of the *Drosophila* humoral response consisting of multiple host defence peptides with specific activities, and demonstrates how natural polymorphisms found in *Drosophila* populations can affect host susceptibility.

## 1.2 Introduction

The ability to rapidly combat pathogens is critical to organism health and survival. Organisms sense natural enemies through pattern recognition receptors, triggering the activation of core immune signalling pathways. These pathways regulate the expression of a plethora of immune effectors that provide a first line of innate defence. It was generally thought that innate immune effectors act together as a cocktail to kill microbes. However recent studies have challenged this view revealing an unexpected high degree of specificity in the effector response to infection [1–3].

Chief amongst immune effectors are antimicrobial peptides (AMPs), host-encoded antibiotics that exhibit microbicidal activities [1,2,4,5]. Insects, and particularly the genetically tractable model *Drosophila*, have been especially fruitful in identifying and characterizing AMP potency and function [4,6–9]. In *Drosophila*, systemic infection triggers the expression of a battery of antimicrobial peptides that are secreted into the hemolymph by the fat body to transform this compartment into a potent microbicidal environment. This systemic AMP response is tightly regulated by two signalling cascades: the Toll and Imd pathways. These two pathways are similar to mammalian TLR and TNFalpha NF-κB signalling that regulate the inflammatory response [10,11]. They are differentially activated by different classes of microbes. The Toll pathway is predominantly instigated after sensing infection by Gram-positive bacteria and fungi, while the Imd pathway is especially responsive to Gram-negative bacteria and some Gram-positive bacteria with DAP-type peptidoglycan [11–13]. The expression of each AMP gene is complex, receiving differential input from either pathway, with most AMPs being at least somewhat co-regulated during the systemic immune response [14–16].

In *Drosophila*, several families of AMPs contribute downstream of Toll and Imd. This includes the Cecropin, Attacin, Diptericin, Defensin, Metchnikowin, Drosomycin, Baramicin, and Drosocin gene families [1,3,4]. Other host defence peptide families include Daisho and Bomanin, which are important for defence, but in vitro killing activity is yet to be shown [17,18]. How these immune effectors contribute individually or collectively to host defence remains poorly understood. Use of single and compounds mutants has revealed that defence against some pathogens relies on the collective contributions of multiple AMP families. However recent studies have also shown that single defence peptides can play highly specific and important roles during infection. In one case, *Diptericins* are the critical AMP family for surviving infection by *Providencia rettgeri* bacteria. This specificity is so remarkable that flies collectively lacking five other AMP gene families nevertheless resist *P. rettgeri* infection like wild-type [6], while even a single amino acid change in one *Diptericin* gene can cause pronounced susceptibility to *P. rettgeri* [19]. Studies on Toll effector genes such as *Bomanins, Daishos*, or *Baramicin A* have also found deletion of single gene families can cause strong susceptibilities against specific fungal species [18,20], or mediate general defences against broad pathogen types [17,21]. Lastly, loss of the gene *Drosocin* causes a specific and pronounced susceptibility to infection by *Enterobacter cloacae* [6], agreeing with Drosocin peptide activity in vitro [22]. Unlike the example with *Diptericins* and *P. rettgeri*, other AMPs also contribute collectively to defence against *E. cloacae* [23].

Many AMP genes encode precursor proteins with multiple peptide products processed by furin cleavage [20]. This was initially shown for the Apidaecin gene of honey bees, which produces nine Apidaecin peptides from a single precursor [24]. *Drosophila* also encodes many AMPs with polypeptide precursors. Examples include AMPs of the Attacin and Diptericin gene families [25,26] or *Baramicin A* which encodes three kinds of unique peptide products on a single precursor protein [20,27,28]. Meanwhile, the precursor protein of the nematode AMP “NLP29” is cleaved into six similar Glycine-rich peptides [29,30]. To our knowledge, the independent contributions of sub-peptides from a polypeptide AMP gene has so far never been addressed.

In this study, we reveal that the gene *CG10816* encodes not only the antibacterial Drosocin peptide, but also another host defence peptide produced by furin cleavage of the Drosocin precursor protein. We name this peptide Buletin, and show that it corresponds to IM7, an inducible peptide first identified in 1998 by MALDI-TOF analysis whose gene counterpart was never identified [31]. Using a new mutation affecting only the Drosocin peptide and not Buletin, we show that these two peptides contribute independently to defence against different microbes. Survival analyses show that while Drosocin specifically affects defence against *E. cloacae*, Buletin specifically affects defence against *Providencia burhodogranariea*. Moreover, a previously identified polymorphic site in Buletin (Thr52Ala described in [32]) mirrors the susceptibility effect of Buletin deletion to *P. burhodogranariea*. We therefore uncover a striking example where an AMP-encoding gene produces two peptides with distinct activities. The *CG10816/Drosocin* gene is also an example of how an AMP polymorphism can significantly affect the host defence against a specific microbe. Alongside recent findings using Diptericin and *P. rettgeri*, our results highlight how AMP evolution is likely driven by differential activity against ecologically-relevant microbes.

## Results

For clarity of discussion: we will use the shorthand Drc (with a “c”, no italics) to refer to the mature Drosocin peptide. Whenever possible, we will use *CG10816* to refer to the *Drosocin* gene (common shorthand *Dro*, with an “o”, italicized).

### The Drosocin gene CG10816 encodes IM7

Previous proteomic analyses of hemolymph from infected *Drosophila* revealed several Immune-induced Molecules (IMs) [31]. These molecules were annotated as IM1-IM24 according to their mass, and over time each of these IMs was associated with a host defence peptide gene [17,18,20,33]. At this point, only one of the 24 original IMs remains unknown: IM7. Previous efforts were unable to link this 2307 Da peptide to a gene in the *Drosophila* reference genome. However during our studies, we noticed that IM7 was absent in flies lacking 14 AMP genes, indicating that it is likely produced by one of these genes [6,23]. We repeated these MALDI-TOF proteomic experiments with hemolymph samples from flies carrying systematic combinations of AMP mutations, ultimately honing in on the Drosocin-encoding gene *CG10816/Dro*. Two independent *CG10816/Dro* mutants (*Dro*^*SK4*^ and *Dro-AttAB*^*SK2*^) both lack IM7 in MALDI-TOF peptidomic analysis (Fig. 1).

**Figure 1:**
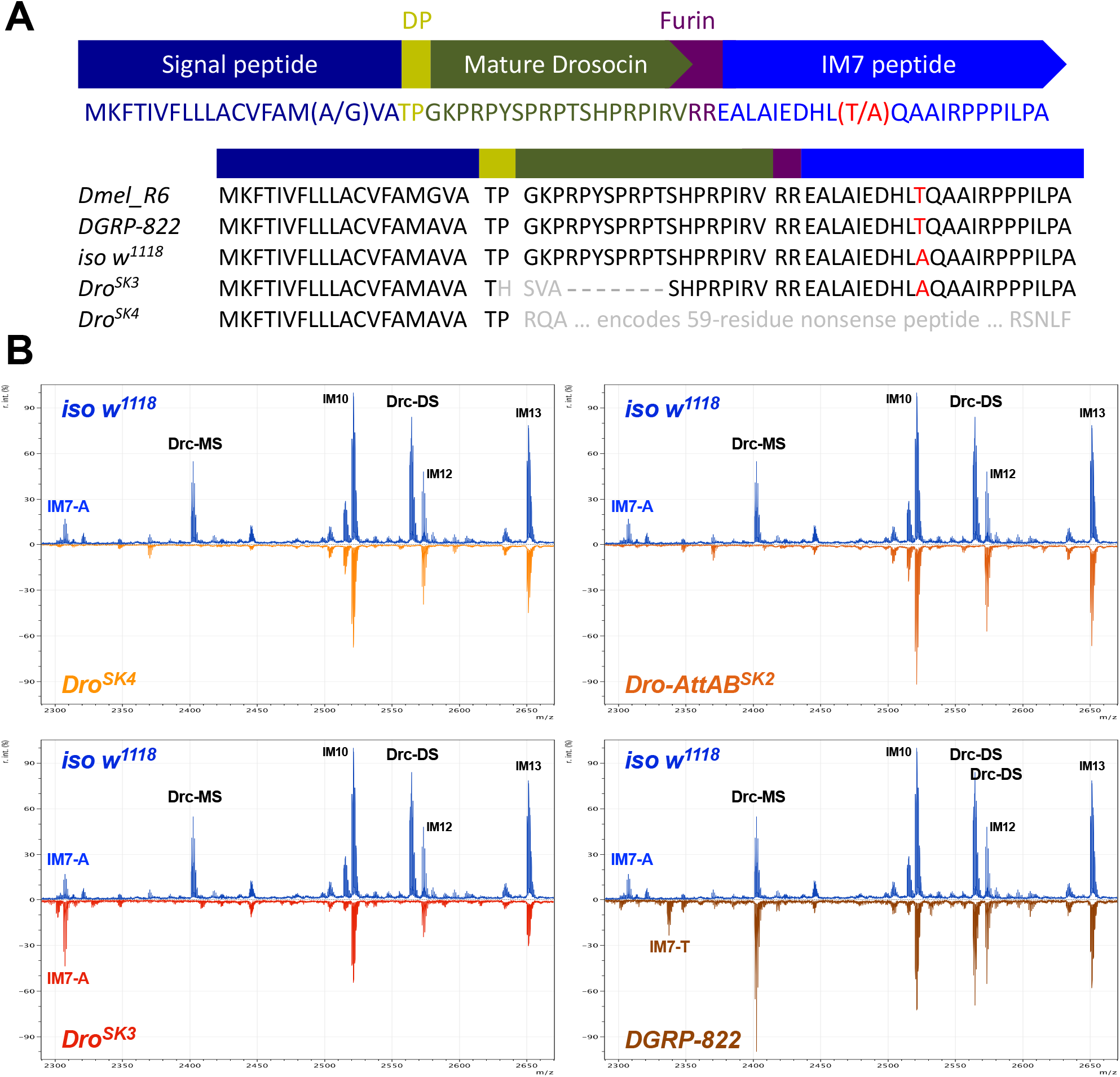
The *CG10816/Dro* gene encodes a polypeptide including both Drc and IM7. A) Overview of the precursor protein structure of the *CG10816/Dro* gene. The Thr52Ala polymorphism in IM7 was noted previously [32]. Here we include an alignment of the CG10816 precursor protein between the Dmel_R6 reference genome and sequences from *iso w*^*1118*^, *Dro*^*SK3*^, *Dro*^*SK4*^, and *DGRP-822* flies. B) MALDI-TOF proteomic data from immune-challenged flies shows that both Drc (Drc-MS, Drc-DS) and the 2307 Da peak of IM7 is absent in *Dro*^*SK4*^ and *Dro-AttAB*^*SK2*^ flies. The frameshift present in *Dro*^*SK3*^ removes the Drc peptide, but does not prevent the secretion of IM7. Threonineencoding IM7 appears in DGRP-822 (2337 Da), alongside loss of the 2307 Da peak.

*CG10816/Dro* was initially identified as a single ORF gene encoding the Drc peptide. Drc is an O-glycosylated Proline-rich peptide that binds bacterial DnaK/Hsp70 similar to other Proline-rich insect AMPs [22,34–36]. Mature Drc requires O-glycosylation for activity, which involves the biochemical linking of either mono-(MS), di-(DS), or rarely tri-saccharide (TS) groups to the Threonine at position 11 of the Drc peptide [22,33]. These different O-glycosylations yield peptides with different mature masses of 2401, 2564, and 2767 Da (Drc-MS, -DS, and -TS respectively). Unmodified Drc peptide has an expected mass of 2199 Da, which is not an intuitive match for the 2307 Da peak of IM7, even considering other post-translational modifications. This suggests that another element of the *CG10816/Dro* gene encodes IM7.

### IM7 is the C-terminus product of the CG10816 precursor

It is puzzling that IM7 could not be annotated to the *CG10816/Dro* gene given that the nucleotide sequence has been known for decades. One previous study noted that the *CG10816/Dro* gene was likely cleaved at a furin-like cleavage site, and had a small undescribed C-terminal peptide [25]. Lazzaro and Clark [32] further described a polymorphism in the *CG10816/Dro* gene encoding either a Threonine or Alanine at residue 52 in the C-terminus of the precursor protein sequence (Thr52Ala). The *D. melanogaster* reference genome encodes the Threonine version of this polymorphism. Using the sequence of the reference genome, the *CG10816* C-terminus mature mass would be 2337 Da without considering post-translational modifications. If we instead substitute an Alanine at this site, the predicted mass of the *CG10816* C-terminus becomes 2307 Da, exactly matching the observed mass of IM7. We confirmed that our wild-type DrosDel isogenic genetic background encodes an Alanine allele both by Sanger sequencing and LC-MS proteomics (data not shown). We next performed MALDI-TOF on the hemolymph of flies from DGRP strain 822 *(DGRP-822)*, which encodes a Threonine in its C-terminus. Exactly matching prediction, *DGRP-822* flies lack the 2307 Da IM7 peak, and instead have a 2337 Da peak that appears after infection (Fig. 1B).

Serendipitously, while generating *CG10816/Dro* mutants using CRISPR-Cas9 we recovered a complex aberrant locus (*Dro*^*SK3*^) that disrupts 11 amino acid residues of the mature Drc peptide, including its critical O-glycosylated Threonine (Fig. 1A). However the *Dro*^*SK3*^ deletion later continues in the same reading frame, including the RVRR furin cleavage site and C-terminus. Thus we suspected that the C-terminal peptide would be secreted normally in *Dro*^*SK3*^ flies. When we ran MALDI-TOF analysis on immune-induced hemolymph from *Dro*^*SK3*^ flies, we recovered a signal that all but confirmed the identity of the *CG10816* C-terminus: *Dro*^*SK3*^ flies lacked the Drc-MS and Drc-DS peaks, but the 2307 Da peak corresponding to IM7 remained immune-inducible (Fig. 1B).

Taken together, we reveal that *CG10816* encodes two peptides: Drc and IM7, which are produced from a precursor protein by cleavage at a canonical furin cleavage site. IM7 is a 22residue peptide with a net anionic charge (−1.9 at pH = 7) that does not share overt similarity with Drc (+5.1 at pH = 7), though both peptides are Proline-rich. A naturally occurring polymorphism previously obscured the annotation of IM7 as a *CG10816* gene product. This analysis was greatly facilitated by the use of newly-available AMP mutations. We name this C-terminal peptide Buletin (Btn) after Philippe Bulet, whose dedicated efforts in the 1980s-1990s characterized many of the *Drosophila* AMPs including *Drosocin* [4,22,37].

### Drc, but not Btn, is responsible for the CG10816-mediated defence against Enterobacter cloacae

Previous studies have suggested that flies lacking just the *CG10816/Dro* locus can resist infection by most bacteria, but are specifically susceptible to infection by *E. cloacae* [6], and also somewhat *E. coli* [38] and *Providencia burhodogranariea* [6]. The fact that *CG10816* encodes not one but two peptides raises the question of the specific contribution of these two peptides to *CG10816* effects. Therefore, we took advantage of the *Dro*^*SK3*^ and *Dro*^*SK4*^ mutations that differently affect the Drc and Btn peptides (Fig. 1A) to explore the respective role(s) these peptides play by comparing the survival of these mutants to different infections. We focused our screen on a panel of Gram-negative bacteria of interest: *E. cloacae β12* bacteria that *CG10816/Dro* mutants are specifically susceptible to [6,23], a recently-isolated *Acetobacter sp*. that can kill AMP mutant flies [39], *E. coli 1106* suggested to be affected by *CG10816/Dro* [22,38], and *P. burhodogranariea strain B* where *CG10816/Dro* was shown to contribute to defence alongside other AMPs [6]. All experiments were performed with wild-type and mutant flies that were isogenized in the DrosDel genetic background according to Ferreira et al. [40].

We found that individual *CG10816/Dro* mutants (both *Dro*^*SK3*^ and *Dro*^*SK4*^) were not overtly susceptible to infection by *E. coli 1106* or *Acetobacter sp. ML04*.*1* (Fig. S1). We could also repeat our previous findings that *Dro*^*SK4*^ and *Dro-AttAB*^*SK2*^ flies were highly susceptible to *E. cloacae* infection, causing 40-50% mortality by 3 days after infection. Importantly, use of *Dro*^*SK3*^ flies that lack Drc but produce Btn confirms that this susceptibility is principally caused by a loss of Drc peptide and not Btn (Fig. 2A): *Dro*^*SK4*^ and *Dro-AttAB*^*SK2*^ flies lacking both Drc and Btn were only slightly more susceptible than *Dro*^*SK3*^ flies lacking Drc alone, a difference that was not statistically significant (*Dro*^*SK4*^ and *Dro-AttAB*^*SK2*^ comparisons to *Dro*^*SK3*^, p > .05 in both cases).

**Figure 2:**
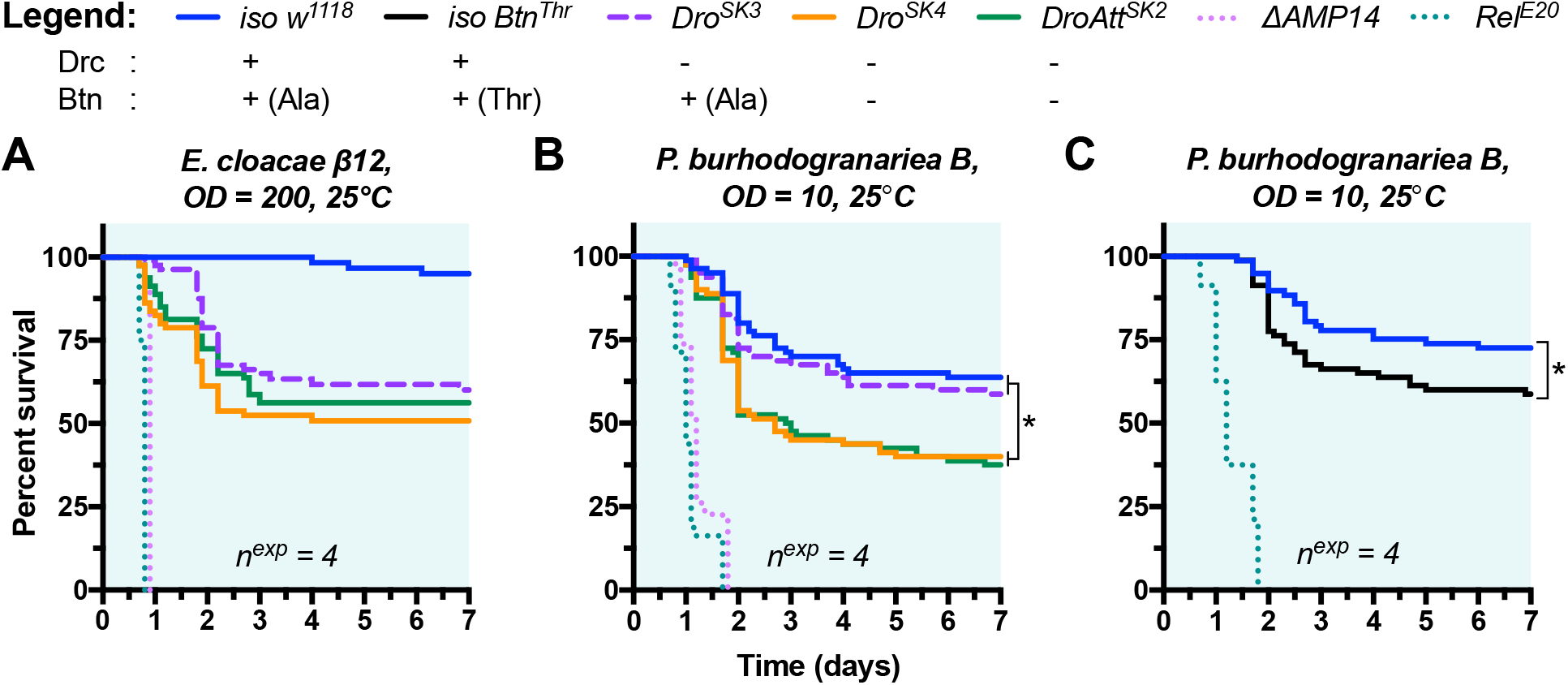
Mutations affecting Buletin cause a specific susceptibility to *P. Burhodogranaria*. A) *Dro*^*SK3*^ flies succumb to infection by *E. cloacae* slightly later than either *Dro*^*SK4*^ or *Dro-AttAB*^*SK2*^ flies that lack both Drc and Btn. The ultimate rate of mortality is comparable (p > .05 in comparisons between these various *Dro* mutants). B) *Drosocin* mutants that retain Btn (*Dro*^*SK3*^) survive infection by *P. burhodogranariea* better than flies lacking both Drc and Btn (*Dro*^*SK4*^, *DroAtt*^*SK2*^). C) Wild-type flies with the Threonine allele of the Btn Thr52Ala polymorphism phenocopy the effect of Btn deletion compared to Alanine-encoding *iso w*^*1118*^ in defence against *P. burhodogranariea*.

Thus, comparison of mutants lacking Drc, or both Drc and Btn, reveals that the *CG10816*mediated defence against *E. cloacae* is specifically mediated by the Drc peptide. Meanwhile Btn does not seem to contribute to defence against this bacterial infection in a significant way.

### Btn, but not Drc, is important for survival to P. burhodogranariea infection

We previously found that *CG10816* could contribute to defence against *P. burhodogranariea* synergistically alongside *Diptericins* and *Attacins* [6]. We next assessed the contribution of our different *CG10816/Dro* gene mutants to defence against *P. burhodogranariea*. To our surprise, the presence or absence of Btn causes a pronounced survival difference after infection by *P. burhodogranariea*: *Dro*^*SK3*^ flies that still produce Btn survive as wild type, while *Dro*^*SK4*^ or *Dro-AttAB*^*SK2*^ flies suffer significantly increased mortality (Fig. 4B). This trend is the opposite of what is observed after infection with *E. cloacae*: Drc does not play an important role in defence against *P. burhodogranariea*, but Btn does. As emphasized by the greater susceptibility of AMP-deficient *ΔAMP14* and Imd-deficient *Rel*^*E20*^ flies (Fig. 2B), Btn deficiency explains only part of the susceptibility to *P. burhodogranariea*. This is consistent with our previous study, which showed that *CG10816/Dro* contributes to defence against this bacterium alongside the contributions of *Diptericin* and *Attacin* genes.

Collectively, our study shows that the *CG10816/Dro* locus encodes two host-defence peptides with distinct activities in vivo. This reinforces the notion that innate immune effectors can have very specific roles in vivo.

### The Thr52Ala polymorphism affects Btn activity against P. burhodogranariea in vivo

The existence of a Threonine/Alanine polymorphic residue in Btn in natural fly populations suggests an arms race between Btn and naturally occurring pathogens. Such polymorphisms are common in AMP genes, and are proposed to reflect host-pathogen coevolutionary selection [41,42]. The *P. burhodogranariea* strain used in this study was originally isolated from the hemolymph of wild-caught flies [43], suggesting it is an ecologically relevant microbe to *D. melanogaster*. This prompted us to investigate the contribution of this polymorphism in defence against *P. burhodogranariea*. We next isolated a Btn-Threonine allele (*Btn*^*Thr*^) that we introgressed into the DrosDel background over seven generations. We infected isogenic *Btn*^*Thr*^ and *Btn*^*Ala*^ (i.e. *iso w*^*1118*^) flies with *P. burhodogranariea* to determine if the Btn polymorphism impacts survival. In these experiments, *iso Btn*^*Thr*^ flies suffered a ∼15% increase in mortality compared to *iso w*^*1118*^ flies with *Btn*^*Ala*^ (Fig. 2C, p = .037). The Cox survival hazard ratio is a measure of effect size. The hazard ratio of *Dro*^*SK4*^ vs. *Dro*^*SK3*^ flies (Fig. 2B) and *iso Btn*^*Thr*^ vs. *iso w*^*1118*^ (Fig. 2C) is nearly-identical (hazard ratios: *Dro*^*SK4*^*-Dro*^*SK3*^ = 0.590, *Btn*^*Thr*^*-iso w*^*1118*^: = 0.584). Thus the effect size of changing the Btn allele from Alanine to Threonine causes the same hazard ratio difference as the effect of Btn deletion.

We therefore uncover an important role of Btn in defence against *P. burhodogranariea*, and reveal that the Btn Thr52Ala polymorphism impacts survival against this ecologically relevant pathogen. Alongside the effect of a polymorphism in Diptericin on survival to *P. rettgeri* [19], here we provide a second example of how a polymorphic residue in an AMP gene significantly impacts survival.

## Discussion

Here we show that the *CG10816/Dro* gene encodes two peptides with distinct activities in vivo. Buletin was not annotated previously due to a polymorphism that masked the identity of this second peptide. Most immune studies have used *Drosophila* strains that encode the *Btn*^*Ala*^ allele (e.g. *Oregon-R* [31], *w*^*1118*^ [44], *DrosDel* [6], or *Canton-S* [45]), while the *D. melanogaster* reference genome encodes the *Btn*^*Thr*^ allele. The gene *CG10816* produces a precursor protein cleaved in two locations: i) after the signal peptide at a two-residue dipeptidyl peptidase site that is nibbled off of the N-terminus of mature Drc (Fig. S3, similar sites noted in [20,46]), and ii) at a furin cleavage motif that separates the Drc and Btn peptides (“RVRR” in *CG10816*). Both cleavage motifs are common in AMP genes, including *Drosophila* Attacins, Defensins, Diptericins, and Baramicins, which all encode mature peptides separated by furin cleavage sites [1,20,25].

The *CG10816/Dro* gene is restricted to the genus *Drosophila* [47]. However phylogenetic inference for AMPs is difficult due to their short size [48,49], and functional analogues of the Drc peptide that may share an evolutionary history are described in many holometabolous insects [50]. It is therefore noteworthy that the range of Buletin is far more restricted: Buletinlike peptides are found only in *Dro* genes of fruit flies ranging from the Melanogaster to Obscura groups, and not in outgroup *Drosophila* species (Fig. S2). The Buletin peptide is therefore an evolutionary novelty of the *CG10816* gene C-terminus. The Thr52Ala polymorphism in Buletin is likely maintained by balancing selection [42], similar to a trade-off between alternate alleles of *Diptericin* in defence against the related bacterium *Providencia rettgeri* [19]. The apparent cost of the Thr52Ala polymorphism to surviving infection by *P. burhodogranariea* suggests an evolutionary trade-off between defence against this bacterium and some other function.

The Drc and Btn peptides are not homologous, although both are rich in Proline residues. However Drosocin is O-glycosylated and has a strong cationic charge (+5.1 at pH = 7), while Buletin is unmodified and has a net anionic charge (−1.9 at pH = 7). AlphaFold predicts Buletin to have an α-helical structure [51]. We screened for Buletin activity in vitro diluted in LB according to Wiegand et al. [52]. However in our conditions, we found no effect of Buletin using either Btn^Thr^ or Btn^Ala^ against *P. burhodogranariea* or *E. coli*, even when co-incubated with sublethal concentrations of Cecropin (Sigma) (Fig. S4). It is possible that Buletin contributes to host defence alongside a co-factor, or protects the host from a virulence factor secreted by *P. burhodogranariea*. We do not wish to rule out a direct action of Btn on bacteria though, as our in vitro conditions could have been sub-optimal for revealing an antimicrobial effect. For instance, an anionic AMP of the Greater wax moth synergizes with Lysozyme to kill *E. coli* [53], and AMPs can act synergistically in vitro through complimentary mechanisms of action [26,36,54,55]. While in vitro approaches are a powerful demonstration for AMP function, we are realizing more and more that this is not sufficient to understand peptide activity in vivo. For example, the activity of azithromycin antibiotic changes 64-fold if tested in standard in vitro conditions or with the addition of human serum [56]. Likewise Bomanin peptides do not display activity in vitro, but Bomanin-deficient hemolymph loses Candida-killing activity [21]. While AMPs were first identified for their potent microbicidal activity in vitro [4,9,57], recent studies in *Drosophila* have recovered striking specificity of AMPs in defence in vivo that was never predicted from in vitro analyses [6,18,19]. These results suggest both in vitro and in vivo approaches are necessary to shed light on host defence peptide activity.

It is striking that the Threonine/Alanine polymorphism in Buletin affects the fly defence against *P. burhodogranariea*. This polymorphism is found in wild populations of *D. melanogaster*, and at high frequencies in the Drosophila Genetic Reference Panel: 29% Threonine, 64% Alanine, 7% unknown at DGRP allele 2R_10633648_SNP [32,58]. A polymorphism in *Diptericin A* causes a profound susceptibility to defence against *Providencia rettgeri* [19], and similar polymorphisms are found in various AMP genes of flies [41,42] and other AMP genes from animals including fish, birds, and humans [59–61]. We now add our study on Buletin and *P. burhodogranariea* to the building evidence that such polymorphisms can have major impacts on microbial control. The existence of polymorphisms in AMP genes could have important implications on the survival of species. For instance: we might wonder if inbreeding in honeybees could have fixed disadvantageous AMP alleles contributing to colony collapse disorder [62]. Reduced AMP expression is also associated with conditions like psoriasis [63] or susceptibility to *Pseudomonas aeruginosa* infections in cystic fibrosis patients [64,65]. A targeted screen has even suggested polymorphisms in human *ß-Defensins* correlate with atopic dermatitis [66]. Could polymorphisms in human AMPs help explain predisposition to similar infectious syndromes?

## Conclusion

By uncovering a novel host defence peptide, our study contributes to a growing body of literature establishing the *Drosophila* systemic infection model as boasting the unique ability to reveal specific interplay of host effector-pathogen interactions. This mode of infection allows the use of the fly hemolymph as an arena to monitor pathogen growth in the presence of effectors, with fly survival as a rapid readout. While previous studies in vitro have suggested fly AMPs had generalist activities, use of specific mutations affecting individual AMP genes has now revealed specific relationships between host and pathogen. Early in vitro studies would never have predicted the highly specific requirement for only single peptides in defence against specific pathogens. Taking lessons from the fly, it should be of significant interest to characterize the differential activity of AMP polymorphisms in humans and other animals, which could reveal critical risk factors for infectious diseases.

## 1.3 Materials and Methods

### Fly genetics

Genetic variants were isogenized into the DrosDel isogenic background over 7 generations as described in [40]. The specific mutations studied here were sourced as follows: the *Dro*^*SK3*^ mutation was generated by CRISPR-Cas9 via gRNA injection as described in [67]. The *Dro*^*SK3*^ sequence was validated by Sanger sequencing and the nucleotide and translated sequence is shown in Figure S3A. *Dro*^*SK3*^ flies encode a truncated version of the Drc peptide lacking its critical Threonine needed for O-glycosylation, and we could detect variants of this truncated Drc peptide in MALDI-TOF spectra with variable degradation of the N-terminus (Fig. S3A-B). The *Btn*^*Thr*^ allele used in this study was originally detected in *Def*^*SK3*^ flies from Parvy et al. [68] by virtue of mutation-specific MALDI-TOF proteomics while screening for possible source genes of IM7. After isogenization, *iso Btn*^*Thr*^ flies were confirmed to have a wild-type *Defensin* gene by PCR. Sequence comparisons were made using Geneious R10.

### Microbe culturing conditions for infections

Bacteria were grown to mid-log phase shaking at 200rpm in their respective growth media (Luria Bertani, MRS+Mannitol) and temperature conditions, and then pelleted by centrifugation to concentrate microbes. Resulting cultures were diluted to the desired optical density at 600nm (OD) for survival experiments, which is indicated in each figure. The following microbes were grown at 37 °C: *Escherichia coli strain 1106* (LB), *Providencia rettgeri* (LB). The following microbes were grown at 29 °C: *Providencia burhodogranariea* (LB) and *Acetobacter sp. ML04*.*1 (MRS+Mannitol*).

### In vitro antibacterial assays

Both the Btn^Thr^ and Btn^Ala^ versions of the 22-residue IM7 peptide were synthesized by GenicBio to a purity of >95%, and silk moth Cecropin A was provided by Sigma-Aldrich at a purity of ≥97%. Peptide preparations were verified by HPLC. Peptides were dissolved in water, and concentrations verified by a combination of BCA assay and Nanodrop A205 readings alongside a BSA standard curve. We screened Btn for activity against both *P. burhodogranariea* and *E. coli* alone at 100µM-1mM, or at 100µM in combination with serially diluted Cecropin concentrations spanning the Cecropin MIC (10µM-0.1µM). Microbes were allowed to grow to log-growth phase, at which point they were diluted to OD = 0.0005 in LB, and then 80μL of this dilute culture was added to 20μL of water or peptide mix to reach desired concentrations in a 96-well plate. Bacteria-peptide solutions were left overnight at room temperature and checked for growth the next morning, and in one experiment optical density at 600nm was recorded every ten minutes using a TECAN plate reader (Fig. S4).

Using these conditions, we found an MIC for Cecropin A against *E. coli 1106* of ∼1µM, agreeing with previous *E. coli* literature [69]. We found an MIC of Cecropin A against *P. burhodogranariea* of ∼5µM, though even 0.63µM delays growth by ∼3 hours compared to no-peptide controls (Fig. S4). Even at 1mM, neither the Btn^Thr^ nor Btn^Ala^ showed any growth inhibition alone, and 100µM peptide combinations with Cecropin A showed no reduction of MIC over Cecropin A alone. 100µM represents the upper limit of AMP concentration in fly hemolymph after infection [70], and the concentration of Btn in vivo is likely much lower than this based on MALDI-TOF relative peak intensities [6,20,31,33]. As we tested Btn alone at 1mM, and at 100µM Btn + Cecropin across the Cecropin MIC range, we find that at least in our conditions using LB as diluent, Btn does not display in vitro activity.

### Survival experiments

Survival experiments were performed as previously described [6], with 20 flies per vial with total replicate experiment number reported within figures (*n*^*exp*^). ∼5 day old males were used in experiments, pricked in the thorax at the pleural sulcus. Flies were flipped thrice weekly. Statistical analyses were performed using a Cox proportional hazards (CoxPH) model in R 3.6.3.

### Proteomic analyses

Raw hemolymph samples were collected from immune-challenged flies for MALDI-TOF proteomic analysis as described previously [6,31]. In brief, hemolymph was collected by capillary and transferred to 0.1% TFA before addition to acetonitrile universal matrix. Representative spectra are shown. Peaks were identified via corresponding m/z values from previous studies [20,33]. Spectra were visualized using mMass, and figures were additionally prepared using Inkscape v0.92.

## Author contributions

MAH performed bioinformatic analyses and planned and performed infection experiments. BL supervised the project and MAH and BL wrote the manuscript. SK generated and supplied *Dro*^*SK3*^ flies.

## Acknowledgements

This research was supported by Sinergia grant CRSII5_186397 and Novartis Foundation 532114 awarded to Bruno Lemaitre. We would like to thank Adrien Schmid and Jonathan Pittet of the EPFL Proteomics Core Facility (PCF) for their technical expertise.

## Supplementary Figures

**Figure S1:**
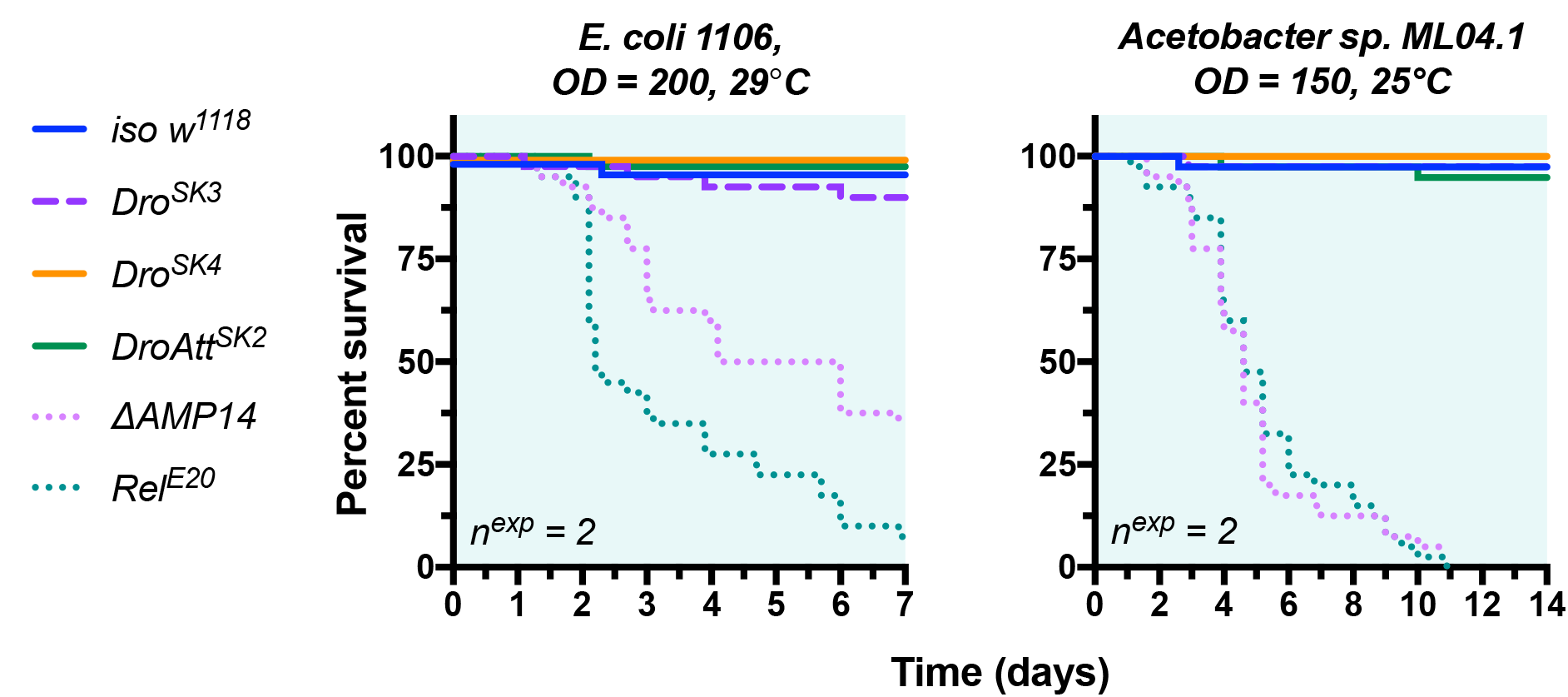
*CG10816/Dro* mutants are not susceptible to *E. coli 1106* or *Acetobacter sp. ML04*.*1* infection. *Rel*^*E20*^ mutants deficient for Imd signalling and *ΔAMP14* flies lacking seven AMP gene families, which includes *CG10816/Dro* deletion, both succumb to these infections, as found previously [6,23,39].

**Figure S2:**
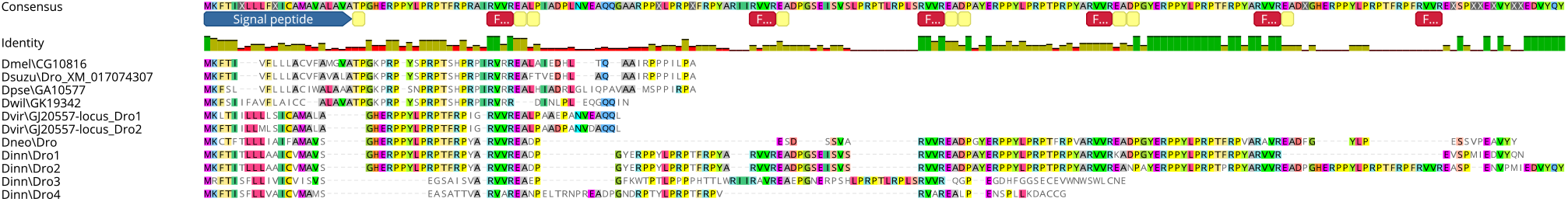
Alignment of Drosocin proteins encoded by various *Drosophila* species. Buletin-like C-terminus peptides are found in *D. pseudoobscura, D. suzukii*, and *D. melanogaster Dro* genes. In *D. willistoni* and subgenus Drosophila flies, Buletin-like peptides are not found. Full precursor protein sequences are shown for each species. Uniquely the *D. neotestacea* and *D. innubila Dro* genes encode multiple Drc peptides in tandem between furin cleavage sites (red boxes at top of alignment) [47]. These furin sites are usually followed by dipeptidyl peptidase sites (yellow boxes at top of alignment), similar to the tandem repeat structure of honeybee *Apidaecin* and *Drosophila Baramicin* [20,24].

**Figure S3:**
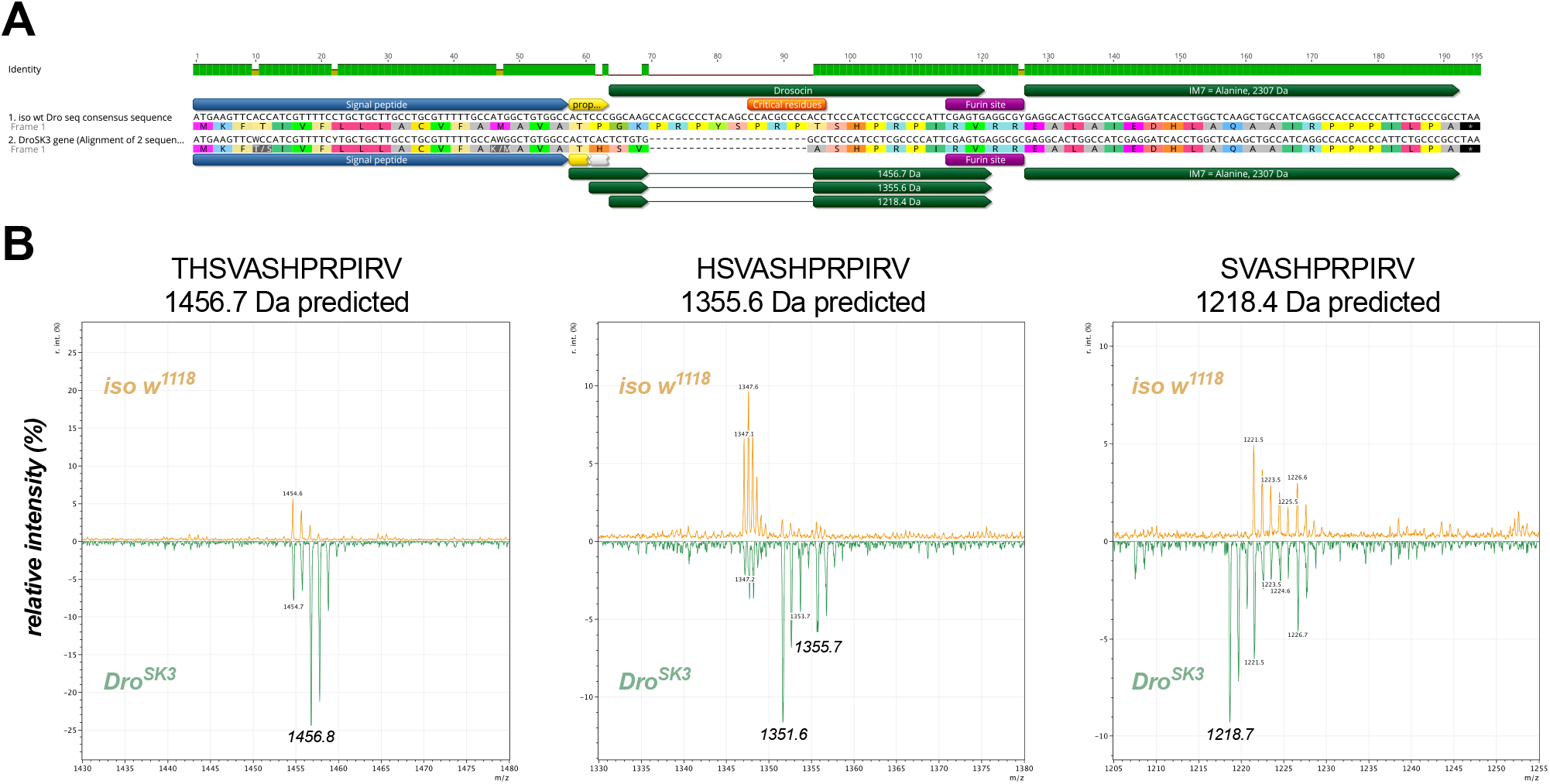
The *Dro*^*SK3*^ mutation deletes the Drc peptide N-terminus, but a truncated Drc peptide is still secreted. A) Alignment and annotation of the nucleotide and mature peptide products of *CG10816* in wild-type and *Dro*^*SK3*^ mutant flies. The *DroSK3* mutation causes a net deletion of 24 nucleotides, and an additional 4 codons (12 nucleotides) are changed. *Dro*^*SK3*^ flies produce Buletin, but also a truncated version of Drc lacking critical residues for activity such as the O-glycosylated Threonine (changed to Alanine, orange critical residues). B) MALDI-TOF spectra show that *Dro*^*SK3*^ flies have unique peptides corresponding to different versions of the *Dro*^*SK3*^ truncated Drc peptide with progressively degraded N-termini (MALDI-TOF reflectron mode). This confirms that a truncated Drc peptide is produced and secreted, though it lacks the critical PRPT motif needed for O-glycosylation. This truncated Drc peptide also lacks dipeptidyl peptidase activity as it is mutated in the CG10816 dipeptidyl peptidase site “TP” ==> “TH” (yellow/grey annotations in A). As a consequence, it is apparently secreted at full length after the signal peptide, only to be progressively degraded from the N-terminus in the hemolymph.

**Figure S4:**
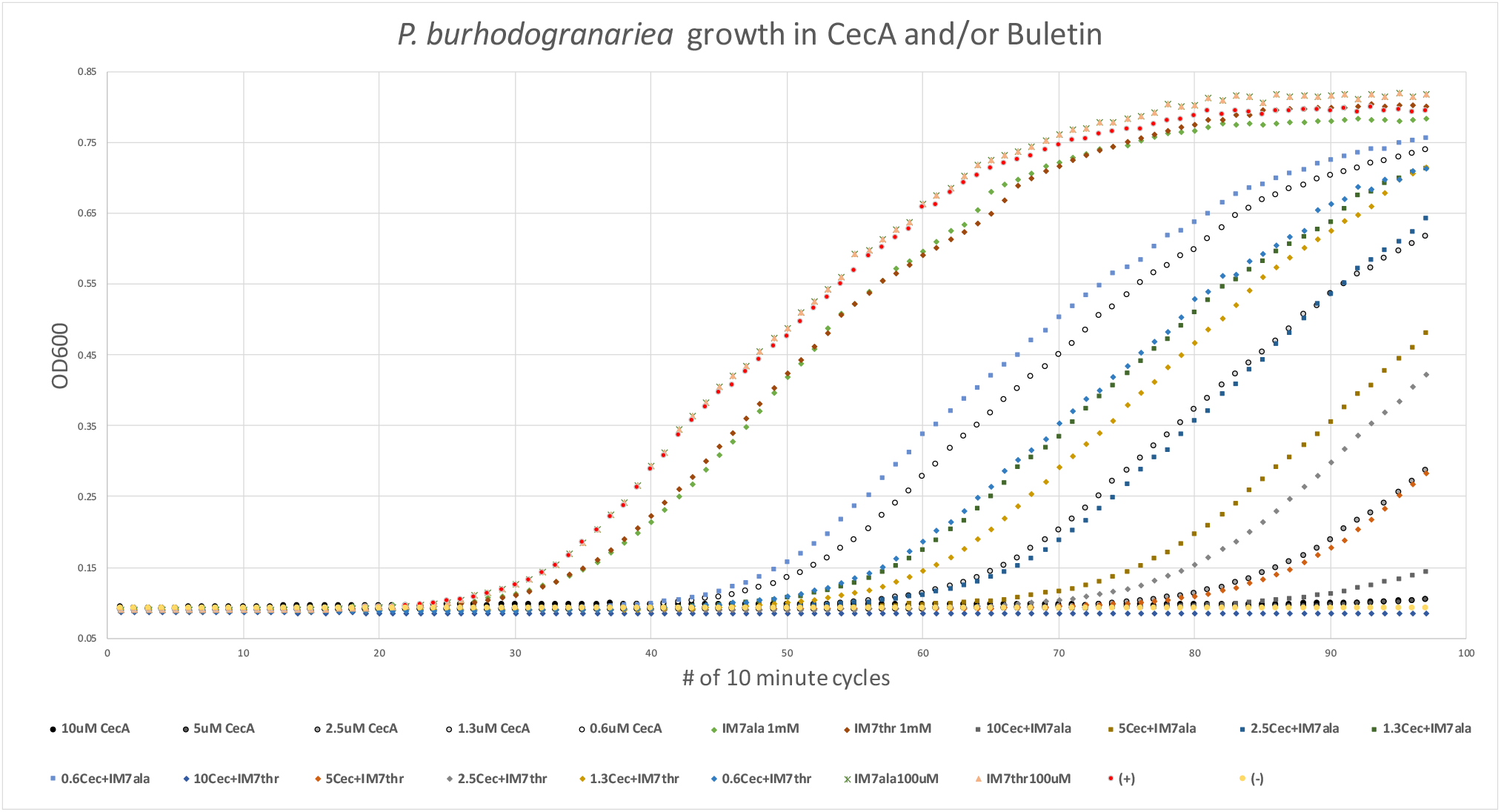
Representative experiment of Cecropin A and Buletin in vitro activity against *P. burhodogranariea*. Bacteria were mixed with peptide in LB and allowed to grow shaking at room temperature overnight. Every 10 minutes, the absorbance at OD600 was recorded. Almost no growth was recorded in the observation period for *P. burhodogranariea* in the presence of 5-10µM Cecropin A. This result is consistent with a separate experiment where we monitored mixtures for bacterial growth only at the end (not shown), suggesting an MIC of Cecropin A against *P. burhodogranariea* of ∼5µM. We conclude that in these in vitro conditions, Buletin does not impact *P. burhodogranariea* growth alone or in combination with pore forming Cecropin peptides.

## Notes

### Competing Interest Statement

The authors have declared no competing interest.

